# The Acute COPD Exacerbation Prediction Tool (ACCEPT): development and external validation study of a personalised prediction model

**DOI:** 10.1101/651901

**Authors:** Amin Adibi, Don D Sin, Abdollah Safari, Kate M Johnson, Shawn D Aaron, J Mark FitzGerald, Mohsen Sadatsafavi

## Abstract

**Background:** Accurate prediction of exacerbation risk enables personalised chronic obstructive pulmonary disease (COPD) care. We developed and validated a generalisable model to predict the individualised rate and severity of COPD exacerbations.

**Methods:** We pooled data from three COPD trials on patients with a history of exacerbations. We developed a mixed-effect model to predict exacerbations over one-year. Severe exacerbations were those requiring inpatient care. Predictors were a history of exacerbations, age, sex, body mass index, smoking status, domiciliary oxygen therapy, lung function, symptom burden, and current medication use. ECLIPSE, a multicentre cohort study, was used for external validation.

**Results:** The development dataset included 2,380 patients (mean 64·7 years, 1373 [57·7%] men, mean exacerbation rate 1·42/year, 0·29/year [20·5%] severe). When validated against all COPD patients in ECLIPSE (n=1819, mean 63·3 years, 1186 [65·2%] men, mean exacerbation rate 1·20/year, 0·27/year [22·2%] severe), the area-under-curve was 0·81 (95%CI 0·79–0·83) for ≥2 exacerbations and 0·77 (95%CI 0·74–0·80) for ≥1 severe exacerbation. Predicted rates were 0·25/year for severe and 1·31/year for all exacerbations, close to the observed rates (0·27/year and 1·20/year, respectively). In ECLIPSE patients with a prior exacerbation history (n=996, mean 63·6 years, 611 (61·3%) men, mean exacerbation rate 1·82/year, 0·40/year [22·0%] severe), the area-under-curve was 0·73 (95%CI 0·70–0·76) for ≥2 exacerbations and 0·74 (95%CI 0·70–0·78) for ≥1 severe exacerbation. Calibration was accurate for severe exacerbations (predicted=0·37/year, observed=0·40/year) and all exacerbations (predicted=1·80/year, observed=1·82/year). The model is accessible at http://resp.core.ubc.ca/ipress/accept.

**Interpretation:** This model can be used as a decision tool to personalise COPD treatment and prevent exacerbations.

**Research in context:** *Evidence before this study:* Preventing future exacerbations is a major goal in COPD care. Because of adverse effects, preventative treatments should be reserved for those at a higher risk of future exacerbations. Predicting exacerbation risk in individual patients can guide these clinical decisions. A 2017 systematic review reported that of the 27 identified COPD exacerbation prediction tools, only two had reported external validation and none was ready for clinical implementation. To find the studies that were published afterwards, we searched PubMed for articles on development and validation of COPD exacerbation prediction since 2015, using the search terms “COPD”, “exacerbation”, “model”, and “validation”. We included studies that reported prediction of either the risk or the rate of exacerbations and excluded studies that did not report external validation. Our literature search revealed two more prediction models neither of which was deemed generalisable due to lack of methodological rigour, or local and limited nature of the data available to investigators.

*Added value of this study:* We used data from three randomised trials to develop ACCEPT, a clinical prediction tool based on routinely available predictors for COPD exacerbations. We externally validated ACCEPT in a large, multinational prospective cohort. To our knowledge, ACCEPT is the first COPD exacerbation prediction tool that jointly estimates the individualised rate and severity of exacerbations. Successful external validation of ACCEPT showed that its generalisability can be expanded across geography and beyond the setting of therapeutic trials. ACCEPT is designed to be easily applicable in clinical practice and is readily accessible as a web application.

*Implications of all the available evidence:* Current guidelines rely on a history of exacerbations as the sole predictor of future exacerbations. Simple clinical and demographic variables, in aggregate, can be used to predict COPD exacerbations with improved accuracy. ACCEPT enables a more personalised approach to treatment based on routinely collected clinical data by allowing clinicians to objectively differentiate risk profiles of patients with similar exacerbation history. Care providers and patients can use individualised exacerbation risk estimates to decide on preventive therapies based on objectively-established or patient-specific thresholds for treatment benefit and harm. COPD clinical researchers can use this tool to target enriched populations for enrolment in clinical trials.

## 1. Background and Objectives

Chronic Obstructive Pulmonary Disease (COPD) is characterised by symptoms of breathlessness and cough, which worsen acutely during exacerbations.^1^ COPD is known to be a heterogeneous disorder with large variations in the risk of exacerbation across patients.^2^ In clinical practice, a history of two or more exacerbations and one severe exacerbation per year is used to guide therapeutic choices for exacerbation prevention.^3^ However, this approach is clinically limited owing to significant heterogeneity in risk even within those who frequently exacerbate.^4^

Prognostic clinical prediction tools enable personalised approaches to disease management. Despite potential benefits, no such tool is routinely used in the clinical management of COPD. This is unlike COPD-related mortality for which clinical scoring schemes such as the BODE index are available and frequently used.^5^ A 2017 systematic review by Guerra and colleagues identified 27 prediction tools for COPD exacerbations.^6^ Among these, only two reported on the validation of the model and none were deemed ready for personalised COPD management in the clinic.^6^

Here, we describe a new model, the Acute COPD Exacerbation Prediction Tool (ACCEPT), to predict, at an individual level, the rate and severity of COPD exacerbation and report on its performance in an independent external cohort, and explain, using case studies, its potential clinical application. As a decision tool, ACCEPT provides a personalised risk profile that allows clinicians to tailor treatment regimens to the individual needs of the patients.

## 2. Methods

In reporting our prediction model, we have followed recommendations set forth by the Transparent Reporting of a Multivariable Prediction Model for Individual Prognosis or Diagnosis (TRIPOD) Statement.^7^

### Target population and source of data

We developed the model using data from COPD patients, without a prior or current history of asthma, and who had experienced at least one exacerbation over the previous 12 months. We then externally validated the model: 1) in COPD patients regardless of their exacerbation history, and 2) in a subset of COPD patients with at least one exacerbation over the previous 12 months. For discovery, we pooled data across all arms of three randomised controlled trials: Macrolide Azithromycin to Prevent Rapid Worsening of Symptoms in COPD (MACRO)^8^, Simvastatin for the Prevention of Exacerbations in Moderate-to-Severe COPD (STATCOPE)^9^, and the Optimal Therapy of COPD to Prevent Exacerbations and Improve Quality of Life (OPTIMAL)^10^. In a secondary analysis, we only used the placebo arms of the trials. We used Evaluation of COPD Longitudinally to Identify Predictive Surrogate End-Points (ECLIPSE)^11^ – an independent longitudinal COPD cohort study – for external validation. The details of each of these studies have been previously published. Briefly, the MACRO study^8^ evaluated the effect of daily low-dose azithromycin therapy on the rate of exacerbations in COPD patients; the STATCOPE study evaluated the effects of daily simvastatin therapy on the rate of exacerbation^9^, and the OPTIMAL study evaluated the effects of tiotropium-fluticasone-salmeterol on the rate of exacerbation compared with tiotropium-fluticasone and tiotropium alone.^10^ In all three trials, which comprised the development dataset, patients who had a history of at least one exacerbation over the previous 12 months were recruited. ECLIPSE, on the other hand, was a multicentre three-year, non-interventional observational study whose primary aim was to characterise COPD phenotypes and identify novel markers of disease progression.^11^ The ECLIPSE study included patients irrespective of their prior history of an exacerbation. Table 1 provides a detailed summary of these studies.

**Table 1.** Available datasets with data on the rate, time, and severity of COPD exacerbations

| Data Source | Design | Intervention | Study Period (Follow-up) | Centres | Inclusion Criteria | Exclusion Criteria |
| --- | --- | --- | --- | --- | --- | --- |
| ***Development*** |  |  |  |  |  |  |
| MACRO | Randomized trial | Azithromycin | 03–2006 to 07–2010 (1 year) | 17 sites in US | >40 yr. Clinical diagnosis of COPD. >10 pack-years of smoking. O<sub>2</sub> or systemic glucocorticoids therapy in the last yr. Hospitalization or ER visit. | Asthma. Exacerbation in the last month. HR above 100 /min. QT<sub>C</sub> >450ms. QT<sub>C</sub> prolonging or TdP-related medication except for amlodipine. Hearing impairment. |
| STATCOPE | Randomized trial | Simvastatin | 03–2010 to 01–2014 (~2 years) | 45 sites (29 in US and 16 in Canada) | 40–80 yrs. Clinical diagnosis of COPD. >10 pack-years of smoking. O<sub>2</sub> OR systemic glucocorticoids OR antibiotics therapy OR hospitalization OR ER visit in the last yr. | Asthma. Receiving statins, or should have received statins. On drugs that contradicted with statins. Unable to take statins. Active liver disease, alcoholism, or allergy. |
| OPTIMAL | Randomized trial | Tiotropium in combination with Salmeterol or Fluticasone-Salmeterol | 10–2003 to 01–2006 (1 year) | 27 sites in Canada | >35 yr. Clinical diagnosis of COPD. >10 pack-years of smoking. Exacerbation requiring systemic glucocorticoids OR antibiotics therapy in the last yr. | Asthma <40 yrs. CHF with persistent severe LVD. Oral prednisone. Intolerance to tiotropium, salmeterol, or fluticasone-salmeterol. Glaucoma. Urinary tract obstruction. Lung transplant or volume reduction. Diffuse bilateral bronchiectasis. Pregnancy or breastfeeding. |
| ***Validation*** |  |  |  |  |  |  |
| ECLIPSE | Cohort |  | 12–2005 to 02–2010 (3 years) | 46 sites in 12 countries | 40–75 yrs. Clinical diagnosis of COPD. >10 pack-years of smoking. | Respiratory disorders other than COPD. Reported exacerbation in the past month. Significant inflammatory disease. |

### Outcomes

The outcomes of interest were the rates of exacerbations and severe exacerbations over one year. Exacerbations were the primary outcome of all three trials and a major outcome measure of the ECLIPSE study. All studies used a similar definition of exacerbations, which was based on the criteria endorsed by the Global Initiative for Chronic Obstructive Lung Disease (GOLD) scientific committee.^3^ Briefly, an exacerbation was defined as an acute episode of intensified symptoms that required additional therapy.^3^ Mild exacerbations were defined as those that were treated with short-acting bronchodilators. Moderate exacerbations were those that required the institution of systemic corticosteroids and/or antibiotics and severe exacerbations were those that required an emergency department visit or a hospitalisation.^3,8–10^

### Predictors

To minimize the risk of bias, optimism, and overfitting, no data-driven variable selection was performed. We pre-specified predictors based on clinical relevance and the availability of predictors in all the datasets. Predictors included non-severe as well as severe exacerbations over the previous year, baseline age, sex, smoking status, percent predicted post-bronchodilator forced expiratory volume in one second (FEV1 % predicted^12^), St. George’s Respiratory Questionnaire (SGRQ) score, body mass index (BMI), and the use of COPD and non-COPD medications as well as domiciliary oxygen therapy during the previous 12 months. COPD medications were defined as long-acting muscarinic receptor antagonists (LAMAs), long-acting β2 agonists (LABAs), and inhaled corticosteroids (ICS). In addition to baseline medications, the model adjusted for treatment assignment in the therapeutic trials (azithromycin in MACRO; statins in STATCOPE; LABA/LAMA and ICS in OPTIMAL). To facilitate clinical implementation, a web application was created (based on conversion factors that have been previously published), which enables the use of a COPD Assessment Test (CAT) score in lieu of SGRQ.^13^

### Follow up

We applied administrative censoring at the one-year follow-up time for patients who had data beyond this threshold. The decision to limit predictions to one year was made *a priori* based on the assumption that predicting exacerbations over this time frame was relevant for clinical COPD management and that prediction accuracy of the model would decrease substantially beyond one year.

### Statistical analysis methods

We used a joint accelerated failure time and logistic model to characterise the rate and severity of exacerbations. We have previously published the details of this approach elsewhere.^14^ In summary, this framework assigns two random-effect terms to each individual, quantifying each individual’s specific rate of exacerbation and the probability that once an exacerbation occurs, it will be severe (appendix p 3). For each patient, this framework fully specifies the hazard of all exacerbations (including their severity) at any given point in time during follow-up, enabling different predictions such as the probability of having a specific number of total and severe exacerbations during the next twelve months.

Two forms of uncertainty in predictions were quantified: the uncertainty due to the finite sample of the validation set (represented by the 95% confidence interval [95%CI] around the mean of projected values), and uncertainty due to the differences in patients’ specific exacerbation frequency and severity (represented by the 95% prediction interval around the mean, the interval which has a 95% probability to contain a future observation of a patient with the same predictors). Shrinkage methods were not applied given the low risk of bias due to complete pre-specification of the model and the relatively high events per predictor in the development dataset.^15^ Because in this framework, the correlation between the previous and future exacerbations rates is modelled through random-effect terms, a history of exacerbation did not enter the model as a predictor. Instead, a Bayesian approach was employed to model the distribution of future exacerbation rate and severity, given the exacerbation history of an individual subject (appendix p 4). The availability of full exacerbation history in the external validation cohort enabled validation of this approach.

We performed statistical analyses using SAS (version 9.4) and R (version 3.6.1).

### External validation

We used the first year of follow-up data in ECLIPSE to establish an accurate 1-year history of exacerbation for each patient. Next, we used the second year of follow-up to validate the model. The model was validated first in the entire COPD cohort of ECLIPSE (n=1,819), and then in a subset of COPD patients who had at least one exacerbation in the first year of follow-up (n=996). This subset was similar to the population characteristics of the development dataset, while the full ECLIPSE cohort enabled assessment of the generalisability of the model beyond patients with an exacerbation history.

We examined model calibration (the degree to which the predicted and actual risks or rates of exacerbations aligned) and discrimination (the extent to which the model separated individuals with different risks).^16^ Calibration was assessed by comparing the predicted and observed exacerbation rates across subgroups with differential risks, evaluating the calibration plots, and calculating Brier scores (i.e. the mean squared error of forecast). Discrimination was assessed by calculating receiver operating characteristic (ROC) curves, and the area-under-the-curve (AUC), and then comparing them using a DeLong’s test.^17^ ROC and AUC calculations were based on the occurrence of two or more exacerbations of any type, or one or more severe exacerbations.^3^

### Ethics approval

The study was approved by the University of British Columbia/Providence Health Research Ethics Board (H11-00786).

## 3. Results

### Participants

Figure 1 presents the flowchart of the sample selection. We excluded 96 patients who were either lost to follow-up (n=33) or had missing values (n=63). The final development dataset included 2,380 patients (1,107 from MACRO, 847 from STATCOPE, and 426 from OPTIMAL; overall mean age 64.7, 1373 [57.7%] males). Patients experienced a total of 3,056 exacerbations, 628 of which were severe. In the external validation dataset, ECLIPSE, 109 patients had missing values. Thus, the final sample included 1,819 COPD patients (mean age 63·3, 1186 [65·2%] male). Among these 996 COPD patients had at least one exacerbation in the first year (mean age 63·6, 611 [61·3%] male). Figure 2 provides a detailed comparison of the development and validation datasets in terms of demographics, predictors, and outcome variables. The average exacerbation rates in the development dataset, the validation set with all patients, and the validation subset containing only those with a prior history of an exacerbation were 1·42, 1·20, and 1·80 events/year, respectively. For severe exacerbations, the average rate was 0·29, 0·27, and 0·40 events/year, respectively.

**Figure 1.**
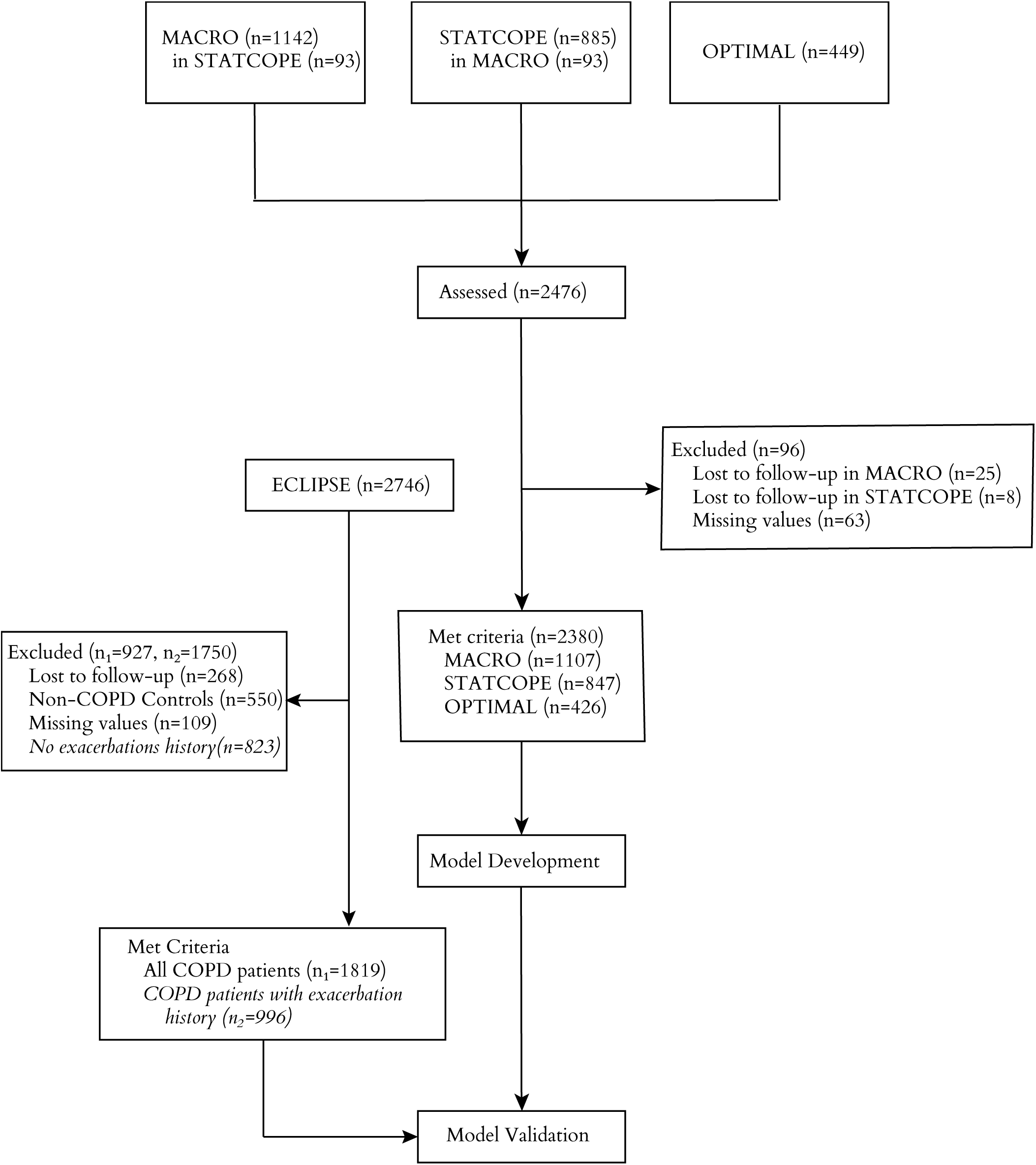
Flow diagram

**Figure 2.**
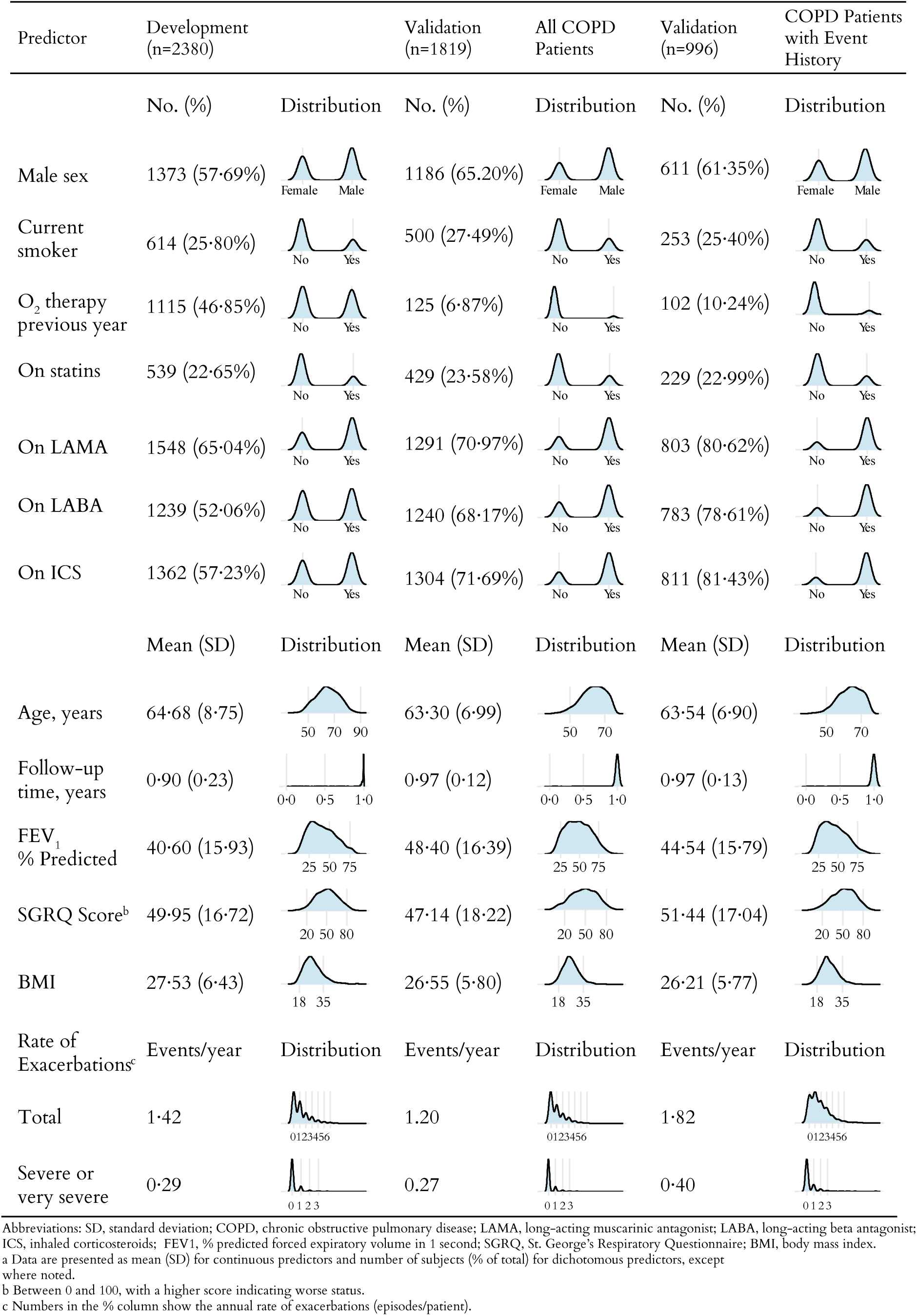
Baseline characteristics

The distribution of baseline predictors among different studies that were included in the development dataset is available in the appendix (pp 5-6). Notably, none of the STATCOPE participants had a history of statin use because patients with cardiovascular comorbidities were excluded from this trial.

We assumed that missing values were missing at random and opted for a complete case analysis given that only 63 out of 2443 patients (2·58%) in the combined development dataset and 109 out of 1928 patients (5·65%) in the validation dataset had missing data (*appendix p 6).*

#### Model specification and performance

Table 2 provides coefficient estimates for predictors. Regression coefficients are shown as log-hazard ratios (ln-HR) for the rate component and log-odds ratios (ln-OR) for the severity component. The full regression results including coefficients representing adjustments for treatments arms are available in the appendix (p 8). Results remained largely unchanged in the secondary analysis based on placebo arms (appendix p 9).

**Table 2.**
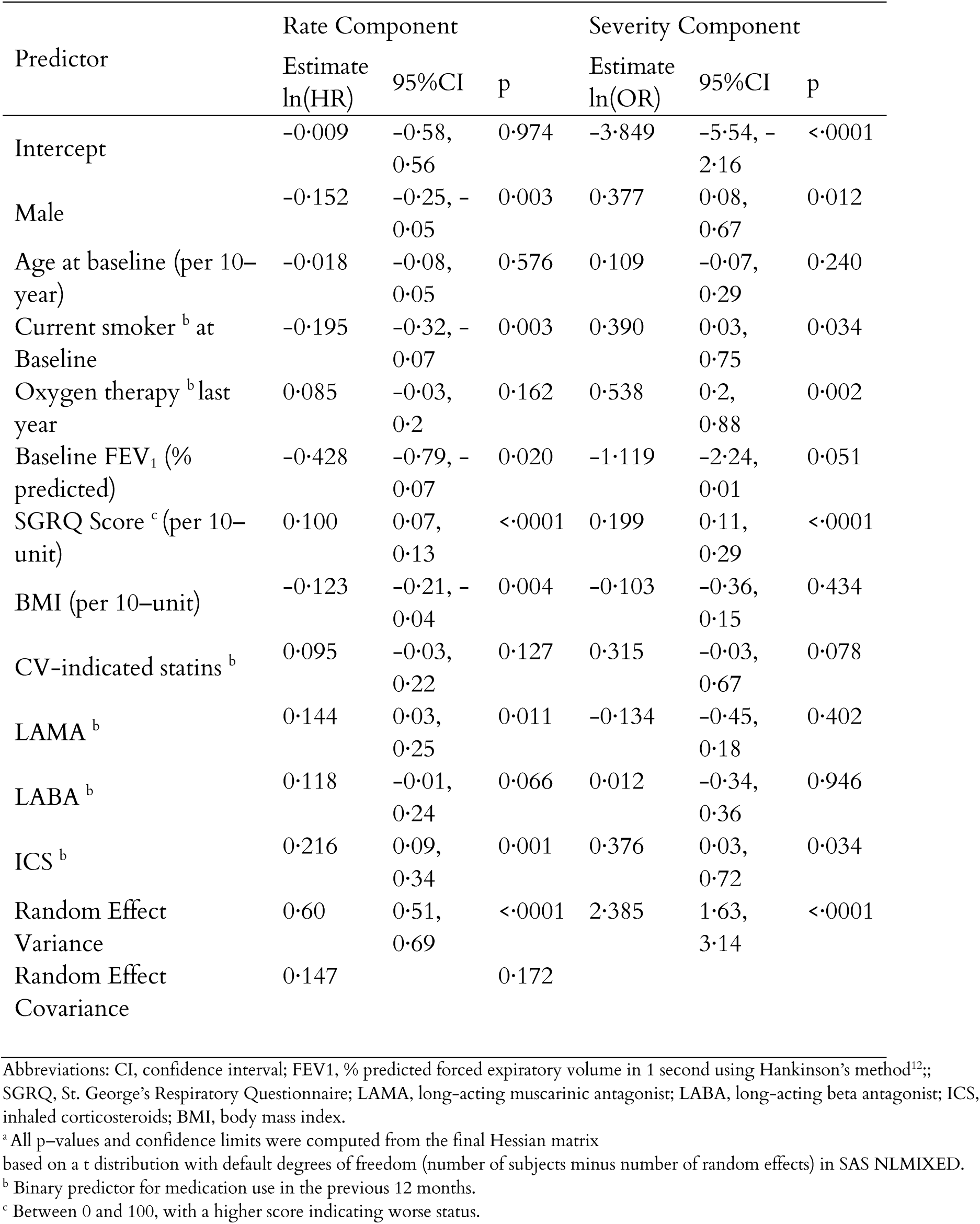
Model coefficients for the joint rate–severity prediction model of COPD exacerbations

When validated against all patients in ECLIPSE (including those without an exacerbation history), ACCEPT slightly overestimated their actual overall exacerbation rates (observed 1·20 events/year, predicted 1·31 events/year) but was accurate for severe exacerbation rates (observed 0·27 events/year, predicted 0·25 events/year, Figure 3*A*). The same trend was observed in all major risk-factor subgroups (Figure 3A) and in both men and women (Figure 4A). The Brier score was 0·20 for all exacerbations and 0·12 for severe exacerbations. In patients with an exacerbation history, ACCEPT showed robust overall calibration: predicted annual exacerbation rate closely matched the observed rate for all exacerbations (observed 1·82 events/year, predicted 1·80 events/year), severe exacerbations (observed 0·40 events/year, predicted 0·37 events/year), and risk-factor subgroups (Figure 3B). Calibration plots comparing per decile average rate of exacerbations showed good agreement between observed and predicted rates for both female and male patients (Figure 4B). The Brier score was 0·17 for all exacerbations and 0·16 for severe exacerbations. Similar results for the development dataset are provided in *the appendix (p 7).*

**Figure 3.**
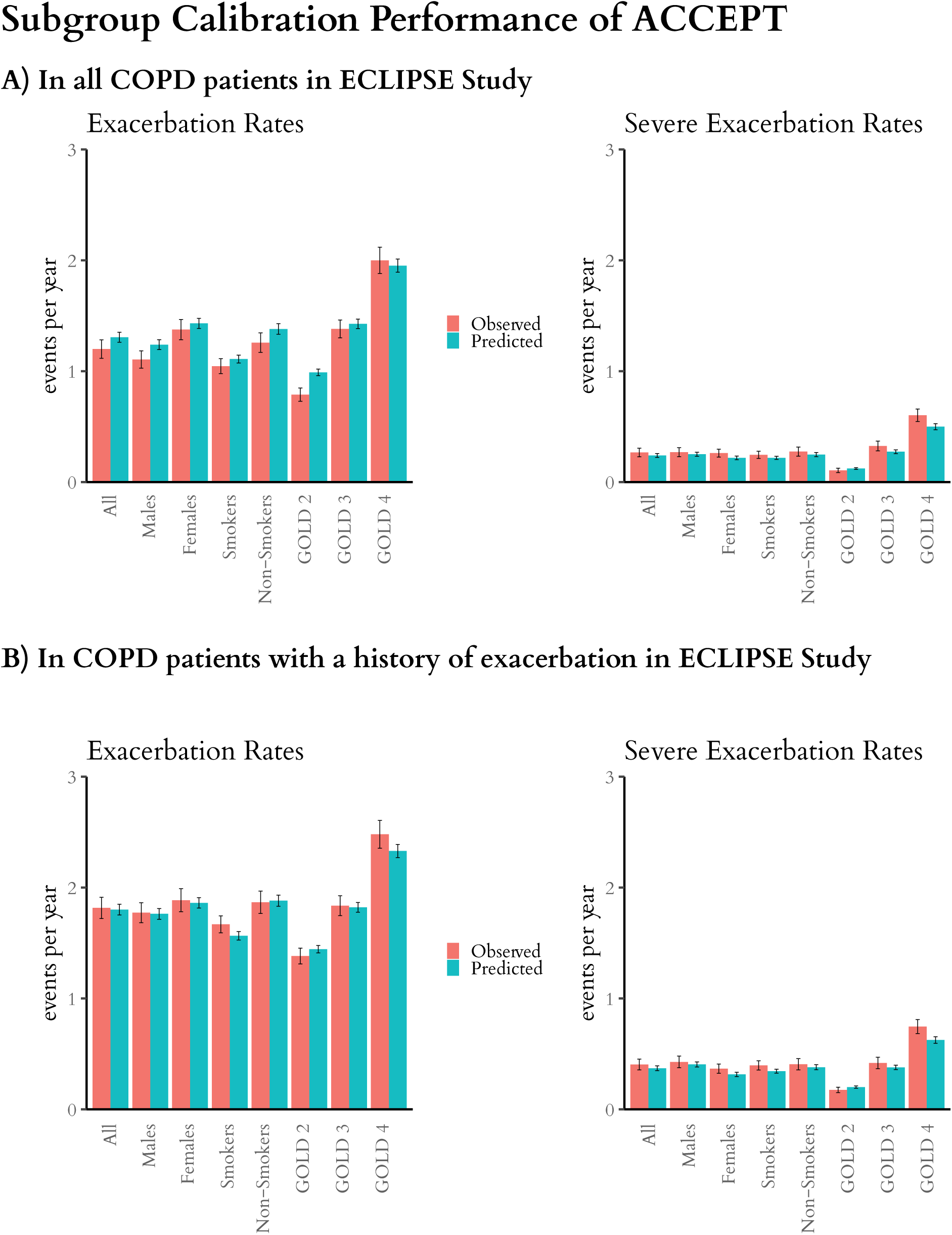
Calibration in risk-factor subgroups

**Figure 4.**
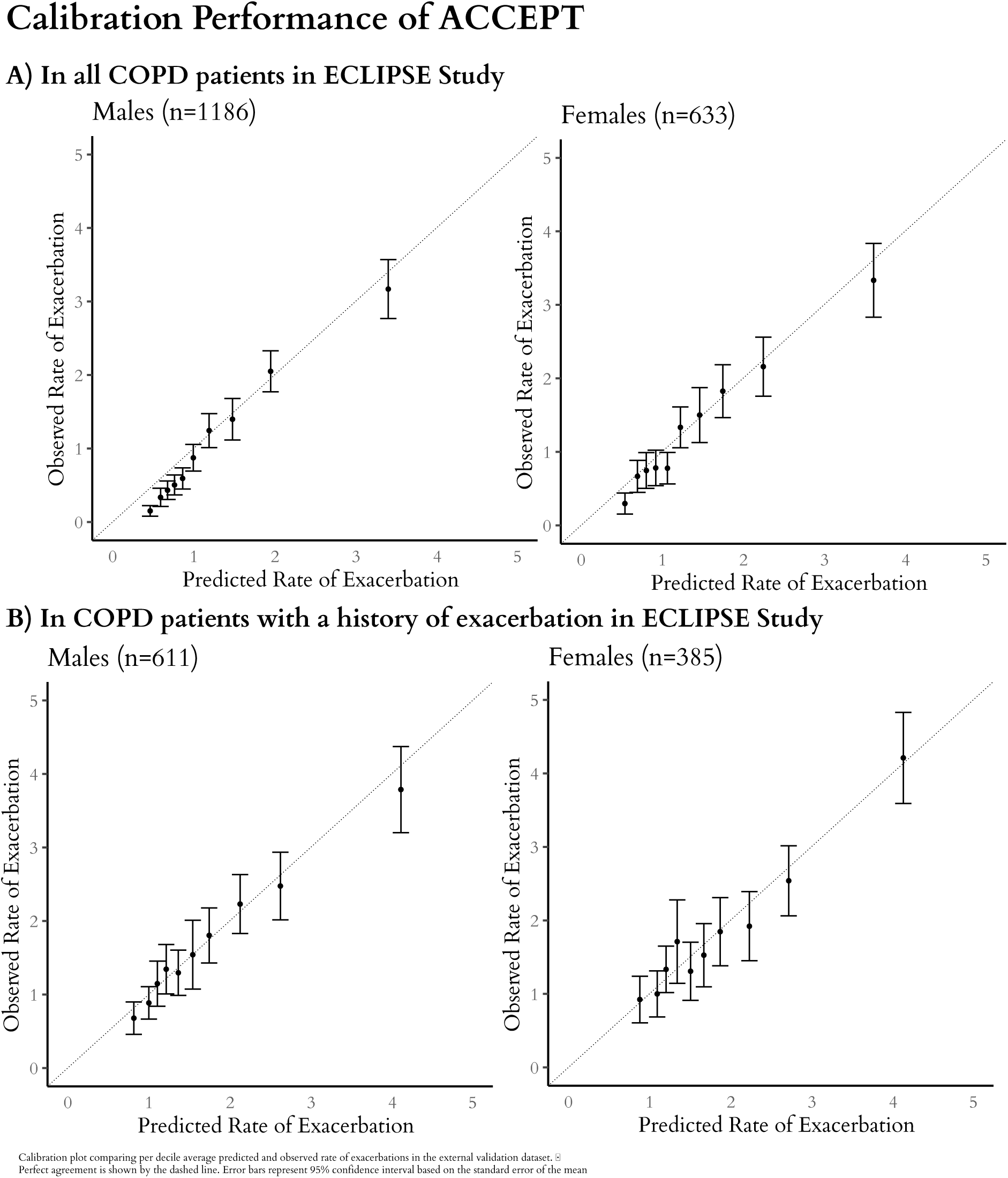
Calibration plot

In all COPD patients, the model had an AUC of 0·81 (95%CI 0·79–0·83) for ≥2 exacerbations and 0·77 (95%CI 0·74–0·80) for ≥1 severe exacerbation (Figure 54*A)*. The corresponding AUCs for COPD patients with an exacerbation history were 0·73 (95%CI 0·70–0·76) for two or more exacerbations and 0·74 (95%CI 0·70–0·78) for at least one severe exacerbation (Figure 5B).

**Figure 5.**
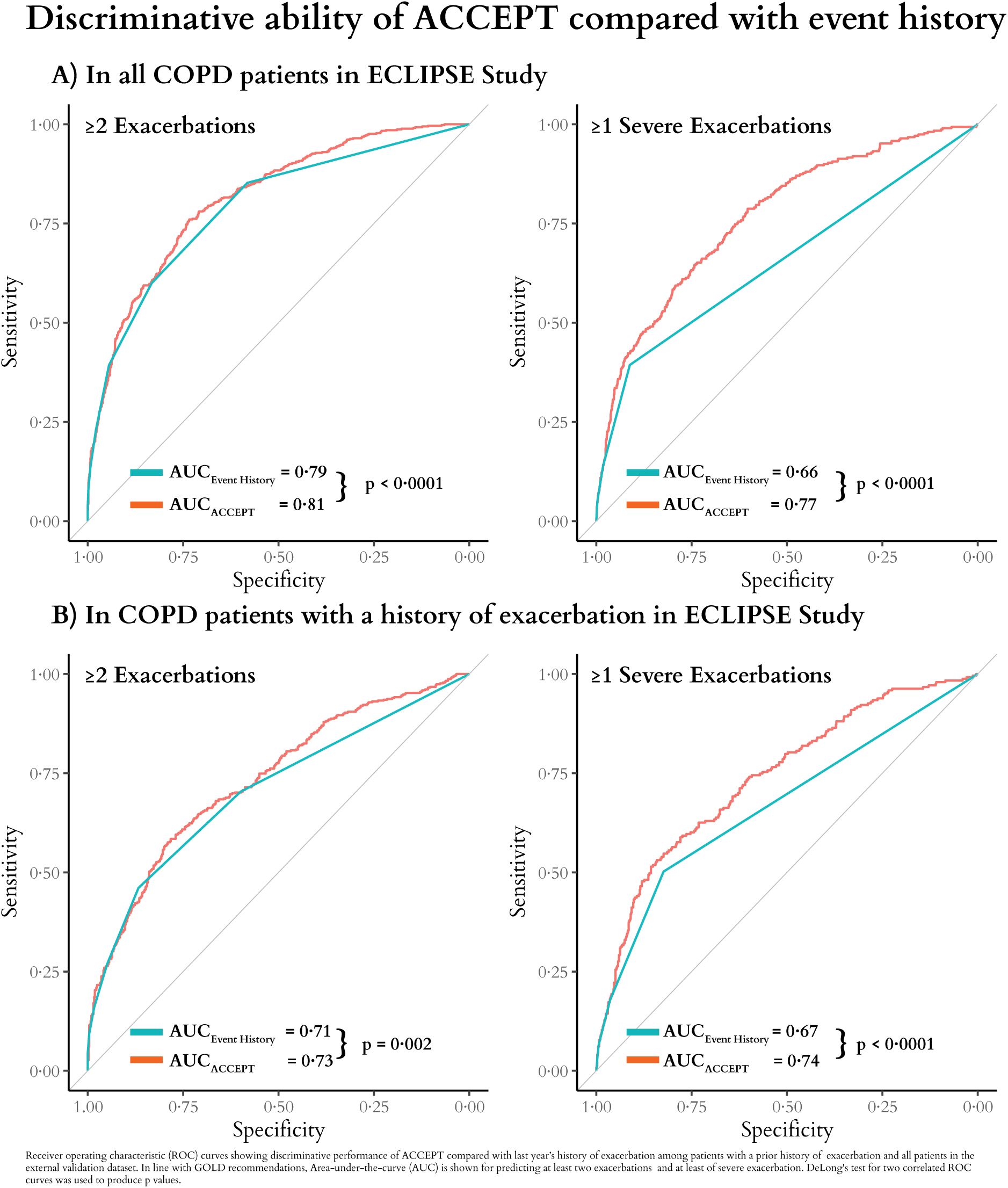
Discriminative ability of ACCEPT compared with event history

Compared to the current practice, which relies exclusively on a prior history of exacerbation to predict future risk of a exacerbation, ACCEPT demonstrated higher performance in predicting severe exacerbations in all COPD patients (AUCACCEPT=0·77 vs. AUCEvent History=0·66, p<0·0001) and in the subset who had a prior history of an exacerbation (AUCACCEPT=0·74 vs. AUCEvent History=0·67, p<0·0001). Similarly, ACCEPT showed better performance for all exacerbations regardless of severity (Figure 5).

## 4. Discussion

The most important finding of the present study was the development and validation of a model (ACCEPT) that uses simple and widely available clinical and demographic variables to predict the risk and severity of exacerbations over a 12-month period, enabling personalisation of care for patients with COPD. In terms of performance, ACCEPT was superior to the current approach of using the individual’s history of exacerbation to predict future risk of exacerbations and in particular for severe exacerbations (where we observed an increase in AUC of 0·11 in all COPD patients and 0·07 in those with an exacerbation in the previous year).

While preventing exacerbations is a major goal in COPD care, there are no tools in practice that can accurately predict the risk or rate of exacerbations in a given individual. Prior studies suggest that patients with a previous history of an exacerbation are more likely to exacerbate in the future than those without.^2^ However, this approach is hampered by a relatively poor resolution, leading to large variations in risk across subjects even among those who have the same history of exacerbations. Our framework builds upon this well-accepted approach and extends its use by incorporating other clinical features that enable a more accurate prediction.

A 2017 systematic review of clinical prediction models for COPD exacerbations found that only two of the 27 reviewed models – CODEX^18^ and Bertens’ model^19^ – reported on any external validation. When the availability of predictors and practical applicability were also taken into account, none of the models were deemed ready for clinical implementation.^6^ We are aware of only two additional prediction models published after this review – by Kerkhof^20^ and Annavarapu^21^ – that have reported external validation. ACCEPT has several notable advantages compared with these models. Importantly, it is externally validated in an independent cohort extending its generalisability beyond therapeutic clinical trials. ACCEPT is also geographically generalisable because the external validation cohort contained data from 12 different countries across North America, Europe, and Oceania. In contrast, previous externally validated models used geographically limited datasets: CODEX was Spanish^18^, Bertens’ model was Dutch^19^, Kerkhof’s model was British^20^, and Annavarapu’s model was based on cross-sectional administrative data from a non-single-payer context in the United States.^21^ Bertens’ model, CODEX, and models by Kerkhof and Annavarapu reported validation AUCs of 0·66 and 0·59, 0·74, and 0·77, respectively. However, the independence of the validation dataset in Kerkhof’s model was questioned as it was selected from the same database as the developmental population. Annavarapu did not report calibration at all, and overall, both models suffered from a lack of generalisability given the local nature of the data that were available to the investigators.

ACCEPT predicts the rate and severity of exacerbations jointly. This is crucial to appropriately tailoring treatments to an individual, as the more granular nature of the output in ACCEPT provides more detailed prediction to assist clinicians in their decision making. For example, ACCEPT can predict the number of exacerbations at a given time period, time to next exacerbation, and the probability of experiencing a specific number of non-severe or severe exacerbations within a given follow-up time (up to one year). This is in contrast to the logistic regression models used in a majority of previous clinical prediction models, which allow prediction probabilities of having at least one exacerbation in a single time window.^6^ Further, this framework can potentially be used for prognostic enrichment of randomised trials by identifying patients who are more likely to exacerbate. Similar to asthma trials, the required sample size and consequently the cost of large trials can be substantially reduced by using prediction models to recruit patients above a certain threshold of expected exacerbation rate.^22,23^

ACCEPT can combine predicted risk with effect estimates from randomised trials to enable personalised treatment. For example, a benefit-harm analysis for roflumilast as preventive therapy for COPD exacerbations reported that the benefits of roflumilast outweighed its potential harm when patients have a severe exacerbation risk of at least 22% over a year.^24^ Using data from this benefit-harm analysis, the accompanying web app of ACCEPT can be used to inform therapeutic decisions on the use of roflumilast for a given patient. Another example is in the potential use of preventative daily azithromycin therapy in COPD. Azithromycin reduces annual exacerbation rate by 27%.^8^ However, it is associated with increased risk of hearing impairment and antimicrobial resistance and thus should be reserved for those at a high risk of future exacerbations.^8^ The accompanying web app illustrates this application by showing the risk of exacerbations with and without daily azithromycin therapy in a given patient. Once care providers discuss the risks of harm and benefits of the therapy and establish patient preference thresholds for the benefit/harm trade-off, ACCEPT can be used to determine whether the preventive azithromycin therapy for that individual reaches or surpasses this threshold.

ACCEPT generates nuanced predictions that allow clinicians to more accurately risk-stratify two patients, who have an identical exacerbation history. The case study in the appendix illustrates this by discussing two patients who have considerably different risk profiles (one projected to experience twice as many severe exacerbations as the other) despite an identical exacerbation history and similar medication profile, smoking status, and age. The complete characteristics of these two patients, as well as prognostic predictions by ACCEPT are available in the appendix (p 2).

### Limitations

The pooled trial data we used to develop the model lacked data on certain variables such as comorbidities, vaccination, blood markers (e.g. eosinophil count), and socio-economic status. As such, these predictors could not be incorporated into the model. Moreover, the developmental dataset did not contain individuals without exacerbations in the previous year; however, the model performed robustly in an external validation dataset that included such patients. Neither the developmental nor the validation datasets included patients with mild (GOLD I) severity and as such, we could not establish the accuracy of predictions for this subgroup. Additionally, our model may not be generalisable to COPD patients with a history of asthma, lifetime non-smokers, patients younger than 40 or older than 80 years of age, or populations outside North America, Europe, and Oceania. Model updating and re-examination of its external validity will be necessary when new sources of data become available.^25^

Compared to simple scoring systems such as the BODE index that can be manually calculated, ACCEPT requires relatively sophisticated computational analysis. While parsimonious models are useful at the bedside, given the complexity of processes involved in the pathogenesis of COPD exacerbations, we believe such tools will have limited resolution. Given the proliferation of hand-held computational devices in clinical practice and the wide availability of clinical parameters that are contained in the model, ACCEPT is usable clinically. This is facilitated through its availability as a web app, a spreadsheet, or the R package ‘accept’.^26^

We emphasise that estimates in our model are predictive and should not be interpreted as “causal”. The observed association between being a smoker and having a lower exacerbation rate (hazard ratio 0·82, 95%CI 0·73–0·93) is one such example. Smoking is likely a marker of disease severity with sicker patients less likely to smoke than those with milder disease. As such, the information in the smoking status variable has high predictive value for the tendency towards exacerbation but is not causally interpretable.

### How to Access the Model

An easy-to-use web application for ACCEPT is available at http://resp.core.ubc.ca/ipress/accept. For any individual patient, the web app predicts 1) the exacerbation risk within the next 12 months, 2) the annual rate of exacerbations, and 3) the probability of experiencing any given number of exacerbations. All outcomes are reported separately for overall and severe exacerbations.

Additionally, we provide an R package^26^, a spreadsheet template, public application programming interfaces (APIs), and additional code at http://resp.core.ubc.ca/research/Specific_Projects/accept.

## 5. Conclusions

ACCEPT is an externally validated and generalisable prediction model that enables nuanced prediction of the rate and severity of exacerbations and provides individualised estimates of risks and uncertainty in predictions. ACCEPT has good to excellent discriminatory power in predicting the rate and severity of COPD exacerbations in all COPD patients and showed robust calibration in individuals with a history of such exacerbations in the past year. Objective prediction of outcomes given each patient’s unique characteristics can help clinicians to tailor treatment of COPD patients based on their individualised prognosis.

## Acknowledgments

We would like to thank Ainsleigh Hill for her contribution to development and documentation of the R package and co-investigators of the Canadian Institutes of Health Research grant Kelly Ablog-Morrant, Drs. Larry Lynd, Teresa To, Annalijn Conklin, Wenjia Chen, Hui Xie, and the Canadian Thoracic Society for their input and feedback.

## Contributors

MS, DDS, JMF, and SDA conceived the study. AA, AS, and MS developed and validated the model. DDS and SDA contributed to data acquisition. AA, KMJ, AS, JMF, DDS, SDA, and MS contributed to the interpretation of the data. AA wrote the first draft of the manuscript and created data visualizations. JMF, DDS, SDA, and MS provided clinical input and oversight. AA developed the Web Application with critical input from KMJ, SDA, DDS, and MS. MS and AA developed the interactive spreadsheet and the R package. All authors revised the manuscript critically and approved the final version to be published.

