## Supplementary Appendix for "The Acute COPD Exacerbation Prediction Tool (ACCEPT): development and external validation study of a personalised prediction model"

**Case Study**

Consider Patient A who has the following characteristics: 57-years of age, male ex-smoker, with a chronic bronchitis phenotype, an $FEV_1$ of 51% of the predicted value (GOLD grade 2), BMI of 18 kg/$m^2$, and a CAT score of 29. Now consider Patient B who is 57 years of age, female ex-smoker with a chronic bronchitis phenotype, an FEV1 of 75% (GOLD 2), a CAT score of 10, and a BMI of 30 kg/$m^2$. Both patients experienced two exacerbations within the past year, one of which was severe, requiring a hospitalisation. Both are using LAMA, LABA, and ICS, but not statins. The complete baseline characteristics of these two patients, as well as their ACCEPT predictions, are shown in Table S1. ACCEPT indicates that Patient A's 1-year probability of experiencing a severe exacerbation is 29% (versus 18% for Patient B). His predicted severe exacerbation rate is 0.6 events/year (versus 0.3 events/year for Patient B). Based on the benefit-harm analysis for roflumilast[1], the web app for ACCEPT predicts that roflumilast is likely to provide a net benefit to Patient A (net benefit probability>80%) but not for Patient B (net benefit probability<20%).

**Table S1 Case study: Despite having the same exacerbation history, predicted severe exacerbation rate and risk for patients A is almost twice as patient B leading to a different recommendation for roflumilast therapy.**

| Predictor | Patient A | Patient B |
|---|---|---|
| Sex | Male | Female |
| Age | 57 | 57 |
| Current smoker | No | No |
| Oxygen therapy last year | No | No |
| $FEV_1$ (% predicted) | 51 (GOLD 2) | 75 (GOLD 2) |
| CAT Score | 29 | 10 |
| BMI | 18 | 30 |
| CV-indicated statins | No | No |
| LAMA | Yes | Yes |
| LABA | Yes | Yes |
| ICS | Yes | Yes |
| Exacerbations in the last year | 2 | 2 |
| Severe Exacerbations in the last year | 1 | 1 |
| **Predictions** | **Patient A** | **Patient B** |
| Predicted Exacerbation Risk | %82 | %79 |
| Predicted Severe Exacerbation Risk | %28 | %18 |
| Predicted Exacerbation Rate | 2.1 | 1.9 |
| Predicted Severe Exacerbation Rate | 0.6 | 0.3 |
| Chance of Net Benefit from Roflumilast | 80%-90% | 10%-20% |
| Roflumilast Prescription | Recommended | Not Recommended |

### Mathematical Definition of the Model

To quantify the incidence and severity of COPD exacerbations and their correlation, we used a parametric joint recurrent-event and logistic regression model similar to the one published previously by our group.[2]

The model consists of two components: a rate component that models the occurrence of exacerbations, and a severity component that models the severity of exacerbations when they occur.

#### The rate component

The rate component was a random-intercept accelerated failure time (AFT) model. Unlike proportional hazard models, AFT models fully specify the likelihood function and as such can be used for prediction of event rates in a new subject. All exacerbations that could occur during follow-up time were considered, with time from baseline to each exacerbation (or censoring) being the unit of analysis. AFT models incorporate the effect of covariates on time-to-events by accelerating or decelerating the passage of time. In such models, the hazard (h) of an event (exacerbation) at time t as a function of the set of covariates (X) is

$h(t) = \theta(X).h_0\big(t.\theta(X)\big)$Where

$$\theta(X) = exp(z_1 + \beta_1.X_1\ +\ \beta_2.X_2\ +\ \dots).$$

The β vector captures the effect of covariates. Between-individual variability (heterogeneity) was modelled through the random-effect term z1. It also captures with within-individual correlation in time to exacerbations.

AFT models require the specification of a baseline hazard ($h_0$). We examined different functions. The function that provided the best fit for the development dataset was Weibull.

#### The severity component

The severity component was a random-intercept logistic regression (binomial distribution with a logit link function). The outcome was the severity of each exacerbation, coded as 1 when the exacerbation was severe, and 0 otherwise.

$$logit\big(P(severity)\big) = exp(z_2 + \beta'_1.X_1\ +\ \beta'_2.X_2\ +\ \dots)$$

Here β' is the vector of coefficients for the severity component, and z2 is the random-effect terms that models individualized risk of an exacerbation being severe, over and beyond the effect of covariates. It also captures within-individual correlation between severity of exacerbations. Individuals contributed to the severity component if they had at least one exacerbation during their follow-up.

The two random-effect terms, z1 and z2, were modelled to have a joint bivariate normal distribution. Any correlation between the rate and severity component (e.g., if frequent exacerbators have a higher proportion of severe to total exacerbations) would be captured in the correlation between z1 and z2, resulting in accurate modelling of dependencies between the two components.

For each individual, this model would generate a predicted time-dependent hazard function for exacerbation, and a predicted risk of an exacerbation being severe. The two quantities can be used to produce a variety of predictions including the number of exacerbations during follow-up, the probability of experiencing any number of exacerbations, the number of severe exacerbations during follow-up, the probability of experiencing any number of severe exacerbations, and so on. The model was coded n PROC NLMIXED in SAS. We used the likelihood-based empirical covariance matrix estimator (otherwise known as the robust or the "sandwich" estimator"). The SAS (version 9.4) code for fitting the model is publicly available at http://resp.core.ubc.ca/research/Specific_Projects/accept.

**Bayesian Recalculation of the Random Effects Distribution to Incorporate Full Exacerbation History**

The joint distribution of random effects for rate and severity of exacerbations were recalculated by giving each pair of random effects the appropriate weight given the observed number of all and severe exacerbations within the past year. We used an iterative process to recalculate random effect distributions. Briefly, let N1 be the number of total exacerbations, and let N2 be the number of severe exacerbations, in the previous year. Let z1 and z2 be two random-effect terms (with bivariate normal distribution). Their joint distribution, P(z1,z2), is estimated in the main model. As well, assuming that the rate of exacerbation does not change between within two consecutive years, the probability of observing a given number of total and severe exacerbations, $P(N1, N2|z1, z2)$, is the main outcome of the model. Applying the Bayes rule. We can calculate the updated distribution of random-effects given a certain exacerbation history:

$$P(z1, z2|N1, N2) \propto P(N1, N2|z1, z2).P(z1, z2)$$

A Monte Carlo simulation with a sample size of 10,000 is used to implement this calculation: first, bivariate random-number generator in R is used to generate a random sample (N=20,000) of z1 and z2. Then the above calculation is performed to assign a weight to each set of (z1, z2) given observed exacerbation history. The weighted (z1, z2) is then used to estimate the distribution of total and severe exacerbations in the next 12 months.

The R code for Bayesian recalculation of random effects distribution is available at ACCEPT's homepage at http://resp.core.ubc.ca/research/Specific_Projects/accept.

### Characteristics of the Development Dataset

#### Figure S1 Baseline characteristics and follow–up statistics in MACRO, STATCOPE, and OPTIMAL [a].

| Variables | MACRO Study (n=1107) | | STATCOPE Study (n=847) | | OPTIMAL Study (n=426) | |
|---|---|---|---|---|---|---|
| | *Distribution* | *No. (%)* | *Distribution* | *No. (%)* | *Distribution* | *No. (%)* |
| Male Sex | 0 1 | 654 (59%) | 0 1 | 478 (56%) | 0 1 | 237 (57%) |
| Current Smokers | 0 1 | 244 (22%) | 0 1 | 256 (30%) | 0 1 | 113 (27%) |
| $O_2$ therapy previous year | 0 1 | 655 (59%) | 0 1 | 408 (41%) | 0 1 | 51 (12%) |
| | *Distribution* | *Mean (SD)* | *Distribution* | *Mean (SD)* | *Distribution* | *Mean (SD)* |
| Age, years | 50 70 90 | 65·18 (8·62) | 50 70 | 62·39 (8·41) | 50 70 90 | 67·89 (8·59) |
| Follow–up time, years | 0.0 0.5 1.0 | 0·93 (0·18) | 0.0 0.5 1.0 | 0·87 (0·25) | 0.0 0.5 1.0 | 0·89 (0·27) |
| $FEV_1$, % predicted | 25 50 75 | 39·55 (15·56) | 25 50 75 | 41·53 (17·65) | 25 50 75 | 41·48 (12·86) |
| SGRQ Score [b] | 20 50 80 | 50·55 (16·4) | 20 50 80 | 49·6 (16·8) | 20 50 80 | 49·1 (17·4) |
| BMI | 18 35 | 26·77 (6·2) | 18 35 | 27·2 (6·9) | 18 35 | 27·5 (6·0) |
| **Exacerbations** | **Frequency** | **Count (Rate [c])** | **Frequency** | **Count (Rate)** | **Frequency** | **Count (Rate)** |
| All | 0123456 | 1597 (1·55) | 0123456 | 850 (1·15) | 0123456 | 596 (1·43) |
| Severe[d] | 0123 | 347 (0·34) | 0123 | 168 (0·23) | 0 1 2 3 | 112 (0·27) |
| | *Distribution* | *No· (%)* | *Distribution* | *No· (%)* | *Distribution* | *No· (%)* |
| Indicated Statin | 0 1 | 446 (40%) | 0 | 0 (0%) | 0 1 | 91 (22%) |
| LAMA | 0 1 | 701 (63%) | 0 1 | 560 (66%) | 0 1 | 282 (67%) |

| | | | | | | |
|---|---|---|---|---|---|---|
| LABA | 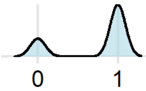 | 822 (74%) | 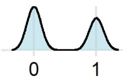 | 359 (42%) | 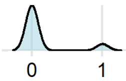 | 53 (13%) |
| ICS | 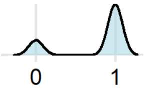 | 856 (77%) | 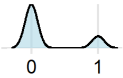 | 181 (21%) | 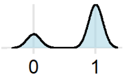 | 317 (76%) |

Abbreviations: COPD, chronic obstructive pulmonary disease; FEV1, forced expiratory volume in 1 second; SD, standard deviation; SGRQ, St. George's Respiratory Questionnaire; LAMA, long acting muscarinic antagonist; LABA, long acting beta antagonist; ICS, inhaled corticosteroids; BMI, body mass index.

[a] Data are presented as mean (SD) for continuous variables and number of subjects (% of column total) for dichotomous variables, except where noted.

[b] Between 0 and 100, with a higher score indicating worse status.

[c] The annual rate of exacerbations (episodes/patient year).

### Table S2 Missing data in the development dataset

| | MACRO (1117 Patients) | OPTIMAL (449 Patients) | STATCOPE (877 Patients) | Total (2443 Patients) |
|---|---|---|---|---|
| ***Complete Cases*** | *1107 (99·10%)* | *426 (94·88%)* | *847 (96·58%)* | *2380 (97·42%)* |
| **Variable** | *Missing N(%)* | *Missing N(%)* | *Missing N(%)* | *Missing N(%)* |
| Male | 0 | 0 | 0 | 0 |
| Age, years | 0 | 7 (1·56%) | 0 | 7 (0·29%) |
| Current Smokers | 1 (0·09%) | 0 | 0 | 1 (0·04%) |
| $O_2$ therapy previous year | 0 | 9 (2·00%) | 0 | 9 (0·37%) |
| $FEV_1$, % predicted | 3 (0·27%) | 1 (0·24%) | 6 (0·68%) | 10 (0·41%) |
| SGRQ Score [b] | 6 (0·54%) | 4 (0·89%) | 24 (2·73%) | 34 (1·39%) |
| BMI | 0 | 2 (0·45%) | 1 (0·11%) | 3 (0·12%) |
| On statins | 0 | 0 | 0 | 0 |
| On LAMA | 0 | 0 | 0 | 0 |
| On LABA | 0 | 0 | 0 | 0 |
| On ICS | 0 | 0 | 0 | 0 |

Abbreviations: COPD, chronic obstructive pulmonary disease; FEV1, % predicted forced expiratory volume in 1 second using Hankinson's method; SGRQ, St. George's Respiratory Questionnaire; ; LAMA, long acting muscarinic antagonist; LABA, long acting beta antagonist; ICS, inhaled corticosteroids; BMI, body mass index.

### Table S3 Missing data in the validation dataset

| | **ECLIPSE (1928 Patients)** |
|---|---|
| ***Complete Cases*** | ***1819 (94·35%)*** |
| **Variable** | ***Missing N(%)*** |
| Male | 0 |
| Age, years | 0 |
| Current Smokers | 16 (0·83%) |
| $O_2$ therapy previous year | 0 |
| $FEV_1$, % predicted | 15 (0·78%) |
| SGRQ Score [b] | 83 (4·30%) |
| BMI | 5 (0·26%) |
| On statins | 0 |
| On LAMA | 0 |
| On LABA | 0 |
| On ICS | 0 |

Distribution of missing values is shown for all COPD patients in ECLIPSE, irrespective of exacerbation history.

Abbreviations: COPD, chronic obstructive pulmonary disease; FEV1, % predicted forced expiratory volume in 1 second using Hankinson's method; SGRQ, St. George's Respiratory Questionnaire; ; LAMA, long acting muscarinic antagonist; LABA, long acting beta antagonist; ICS, inhaled corticosteroids; BMI, body mass index.

### Internal Validation of the Model

The reader can refer to the original publication for a full description of the statistical methodology.[2] The accelerated failure time model requires specification of parametric baseline hazard. We tested exponential, Weibull, log-logistic, and log-normal distributions for the survival function and assessed the internal validity of the resulting models by comparing observed and predicted cumulative number of exacerbations, as well as goodness-of-fit measures of Akaike information criterion (AIC) and Bayesian information criterion (BIC). We selected a Weibull distribution for the survival function because it had the highest agreement between observed and predicted cumulative number of exacerbations (Figure S3) and best goodness-of-fit statistics (Table S4). Figure S2 shows calibration in commonly reported subgroups of sex, smoking status, and disease severity for internal validation. Additionally, the Brier score was 0.21 for all exacerbations and 0.13 for severe exacerbations in the development dataset.

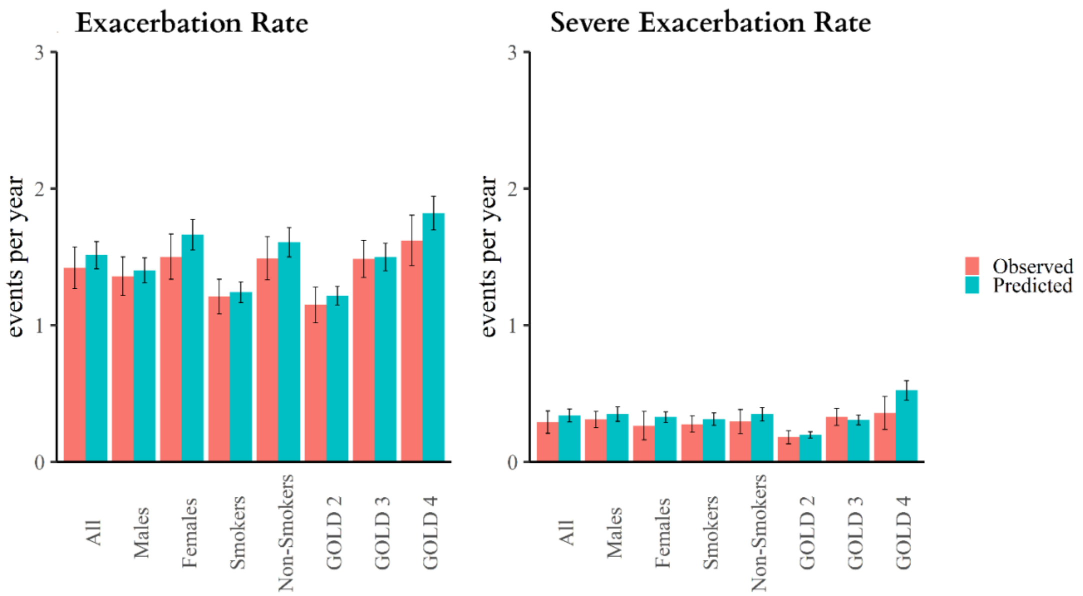

**Figure S2 Calibration of exacerbation and severe exacerbation rate in subgroups of development dataset**

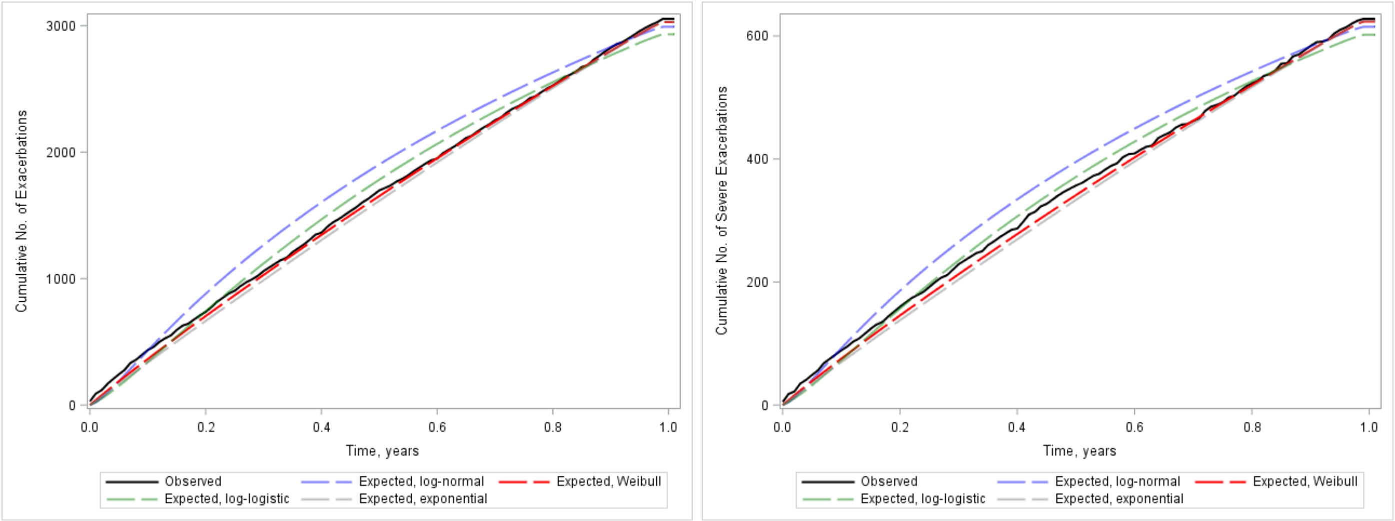

**Figure S3 Internal Validation: Observed and predicted cumulative number of all and severe exacerbations**

**Table S4 Fit statistics for different survival function models.**

| Survival Function Distribution | AIC | BIC |
|---|---|---|
| Weibull Model | 6764·0 | 6995·0 |
| Exponential Model | 6767·1 | 6991·6 |
| Log–Logistics Model | 6821·0 | 7122·0 |
| Log–Normal Mode | 6918·9 | 7219·9 |

Abbreviations: AIC, Akaike information criterion; BIC: Bayesian information criterion.

### Adjustments for Treatment Effects

#### Table S5 Full list of model parameters including adjustments for treatment effects

| Predictor | Rate Component | | Severity Component | |
|---|---|---|---|---|
| | **Estimate ln(HR)** | **Std· Err** | **Estimate ln(OR)** | **Std· Err** |
| Intercept | -0·009 | 0·290 | -3·849 | 0·863 |
| Male | -0·152 | 0·051 | 0·377 | 0·149 |
| Age at baseline (per 10–year) | -0·018 | 0·033 | 0·109 | 0·093 |
| Current smoker at Baseline | -0·195 | 0·066 | 0·390 | 0·184 |
| Oxygen therapy last year | 0·085 | 0·060 | 0·538 | 0·175 |
| Baseline $FEV_1$ (% predicted) | -0·428 | 0·184 | -1·119 | 0·574 |
| SGRQ Score [c] (per 10–unit) | 0·100 | 0·016 | 0·199 | 0·047 |
| BMI (per 10–unit) | -0·123 | 0·043 | -0·103 | 0·131 |
| CV-indicated statins | 0·095 | 0·062 | 0·315 | 0·179 |
| LAMA | 0·144 | 0·056 | -0·134 | 0·160 |
| LABA | 0·118 | 0·064 | 0·012 | 0·177 |
| ICS | 0·216 | 0·064 | 0·376 | 0·177 |
| Randomized Azithromycin | -0·163 | 0·068 | -0·106 | 0·195 |
| Randomized LAMA | 0·167 | 0·111 | 0·104 | 0·337 |
| Randomized LABA | 0·135 | 0·132 | -0·218 | 0·383 |
| Randomized ICS | -0·238 | 0·137 | -0·117 | 0·407 |
| Randomized Statins | -0·054 | 0·079 | 0·106 | 0·229 |
| Random Effect Variance | 0·60 | | 2·385 | |
| Random Effect Covariance | 0·147 | | | |

Coefficients for randomized treatments adjust for treatment effects but are not used for risk prediction.
Abbreviations: CI, confidence interval; FEV1, % predicted forced expiratory volume in 1 second; SGRQ, St. George's Respiratory Questionnaire; LAMA, long-acting muscarinic antagonist; LABA, long-acting beta antagonist; ICS, inhaled corticosteroids; BMI, body mass index.
[a] All p–values and confidence limits were computed from the final Hessian matrix
based on a t distribution with default degrees of freedom (number of subjects minus number of random effects) in SAS NLMIXED.
[b] Binary predictor for medication use in the previous 12 months.
[c] Between 0 and 100, with a higher score indicating worse status.

**Model Based on the Placebo-Group Data**

**Table S6 Model coefficients for the Joint Rate–Severity Prediction Model of COPD Exacerbations Based on Placebo-group data only [a]**

| Predictor | Estimate ln(HR) | p | Estimate ln(OR) | p |
|---|---|---|---|---|
| Intercept | -0·873 | 0·05 | -3·759 | 0·007 |
| Male | -0·230 | 0·003 | 0·537 | 0·03 |
| Age at baseline (per 10–year) | 0·082 | 0·12 | -0·033 | 0·83 |
| Current smoker [b] at Baseline | -0·223 | 0·03 | -0·231 | 0·49 |
| Oxygen therapy [b] last year | 0·057 | 0·53 | 0·320 | 0·23 |
| Baseline $FEV_1$ (% predicted) | -0·260 | 0·34 | -1·468 | 0·10 |
| SGRQ Score [c] (per 10–unit) | 0·146 | <0·0001 | 0·299 | 0·0001 |
| BMI (per 10–unit) | -0·115 | 0·07 | 0·019 | 0·92 |
| CV-indicated statins [b] | 0·084 | 0·37 | 0·424 | 0·13 |
| LAMA [b] | 0·104 | 0·22 | -0·140 | 0·58 |
| LABA [b] | 0·156 | 0·09 | -0·015 | 0·96 |
| ICS [b] | 0·163 | 0·08 | 0·529 | 0·05 |
| Random Effect Variance | 0·563 | <0·0001 | 2·587 | <0·0001 |
| Random Effect Covariance | 0·235 | 0·15 | | |

Abbreviations: CI, confidence interval; FEV1, % predicted forced expiratory volume in 1 second; SGRQ, St. George's Respiratory Questionnaire; LAMA, long-acting muscarinic antagonist; LABA, long-acting beta antagonist; ICS, inhaled corticosteroids; BMI, body mass index.

[a] All p–values and confidence limits were computed from the final Hessian matrix based on a t distribution with default degrees of freedom (number of subjects minus number of random effects) in SAS NLMIXED.

[b] Binary predictor for medication use in the previous 12 months.

[c] Between 0 and 100, with a higher score indicating worse status.

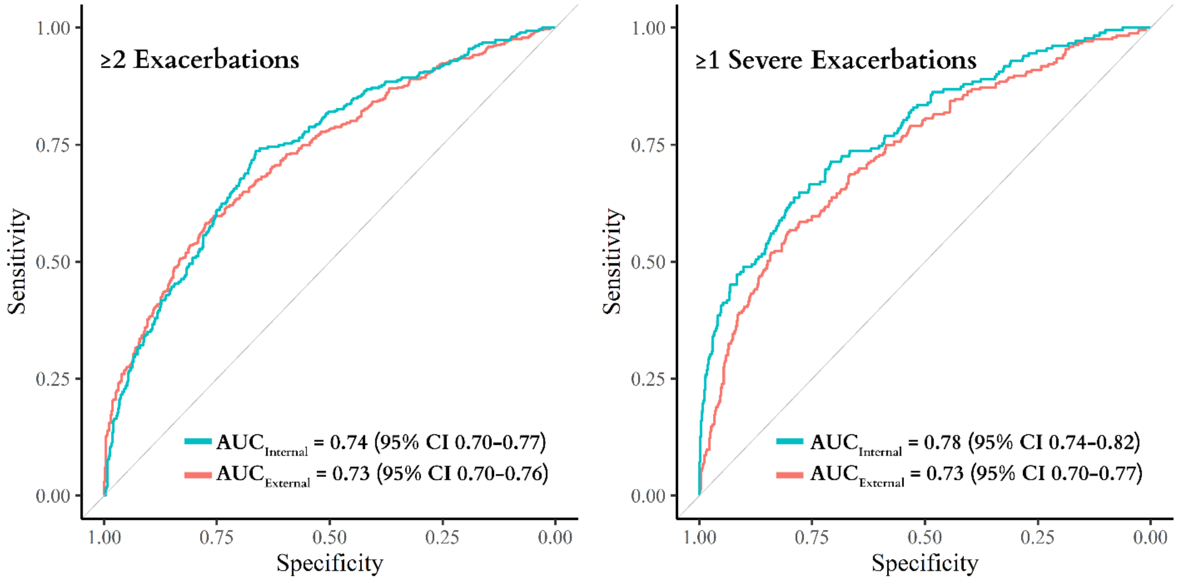

**Figure S4 ROC Curves for a model fitted on placebo arms of the trials only**
